# The Signal in the Noise: Hierarchy and Robustness of Physiological Audience Alignment during Narrative Media

**DOI:** 10.64898/2026.02.05.703516

**Authors:** Ralf Schmälzle, Hee Jung Cho, Monique Turner

## Abstract

Communication research traditionally prioritizes receiver variance, often overlooking how a single message induces commonality across distinct systems. Drawing on classical theory, we propose that effective messages entrain the biological rhythms of the audience. In a large-scale experiment (*N*=198) monitoring eye-tracking, heart rate, and electrodermal activity during a film, we mapped this alignment. Results reveal a processing hierarchy: alignment was strongest for information acquisition (gaze) but also extended to downstream autonomic regulation. Crucially, a temporal manipulation confirmed this state is content-locked: alignment vanished when the narrative sequence was altered but was restored upon computational unscrambling of the physiological time series. This confirms that responses carry a temporal fingerprint of the content. We conclude that messages function as alignment devices, reducing individual noise to create a network of receivers exhibiting shared processing states, thus providing a materially grounded, signal-based definition of the audience.

---

A core tension in communication theory and research lies between the stability of the message signal and the variability of the receiver response. Since the field expanded from early transmission models (Schramm, 1954; Shannon, 1948) and Berlo (1960) formalized the Source-Message-Channel-Receiver (*SMCR*) model, much scholarship has prioritized the final component of this chain: the Receiver. Decades of inquiry examined how individual differences, from demographics to prior knowledge, generate variance in interpretation or other message effects. While this focus on “who” (Lasswell, 1948) is reading, watching, or listening captures the diversity of decoding, it tends to overlook the fundamental mechanism that makes mass communication possible: the ability of a single signal to induce commonality across distinct biological systems. Thus, by focusing on general audience characteristics and varieties of their interpretations, we risk ignoring the baseline convergence in how they process the same signal, the root of the often-invoked “one-to-many” nature of mass communication. We propose a return to the signal, offering a functional definition of the audience: the audience is not a static category of people, but a dynamic state of receiver (*R*) alignment created by the message (*M*) itself.

Theoretically, this biological alignment represents the fidelity of the communication process. In a signal-processing framework, individual receivers are inherently noisy systems; their homeostatic baselines are continuously modulated by idiosyncratic factors such as mood, fatigue, or prior context. For communication to occur, the external signal must be sufficiently structured to overcome this internal noise and induce an aligned state^2^, a phenomenon described in neuroscience as inter-subjective coupling the responses of independent viewers to the temporal flow of the stimulus (Hasson et al., 2004, 2012). Viewed through this lens, a compelling message does not merely “inject” information; rather, it provides a temporal scaffold that coordinates the biological fluctuations of the audience so that the incoming information interacts with receiver-sided decoding processes, but still in a way that should happen similarly across multiple audience members. Consequently, the degree of inter-subjective correlations (ISC) serves as a metric of signal fidelity: the higher the correlation, the more the message has successfully minimized the idiosyncratic noise of individual receivers.

The following article establishes the theoretical rationale for investigating shared processing in mass communication and connects this framework to the measurement of continuous physiological responses while audience members receive and process the same message. We first introduce a theoretical distinction between two layers of alignment: the alignment of information acquisition (visual attention) and that of autonomic regulation (affective resonance). We then outline the basis for shared processing, develop hypotheses regarding the hierarchy and robustness of this alignment, and present evidence that the audience is functionally constituted by the temporal structure of the message itself. After that, we introduce the current study with its hypotheses.

## Theoretical Background

### Shared Processing in Mass Communication

Historically, communication science has struggled to define and study the audience in a way that captures the dynamism of reception (Schramm, 1954). While valuable frameworks exist to describe audiences as publics and interpreters (McQuail, 1997; Schrøder and Gulbrandsen, 2018), these concepts prioritize semantic outcomes. They often overlook the immediate, material reality of the communication process: that distinct receivers are processing the same signal in the exact same temporal order. We propose a shift from treating the audience as a sociodemographic category to viewing it as a functional state of signal-driven alignment. Whether viewers watch a film simultaneously in a theater or asynchronously on a streaming device, they become coupled to the temporal structure of the message (Hasson et al., 2008b; Grall et al., 2021). As illustrated by Schramm’s Tuba model (see Figure 1), mass communication implies the dispersal of identical messages to a multiplicity of receivers. Our approach tests the “neck” of that tuba: the initial moment where the single signal organizes the processing of the many. From this perspective, the audience is a “virtual collective,” brought into existence not by physical co-presence, but by the shared entrainment of their processing states by the message itself. On that communication-theoretical view, a key question is thus not just “who are these people?” or “what are they watching?”, but “to what extent does the message successfully align their processing states?”

**Figure 1.**
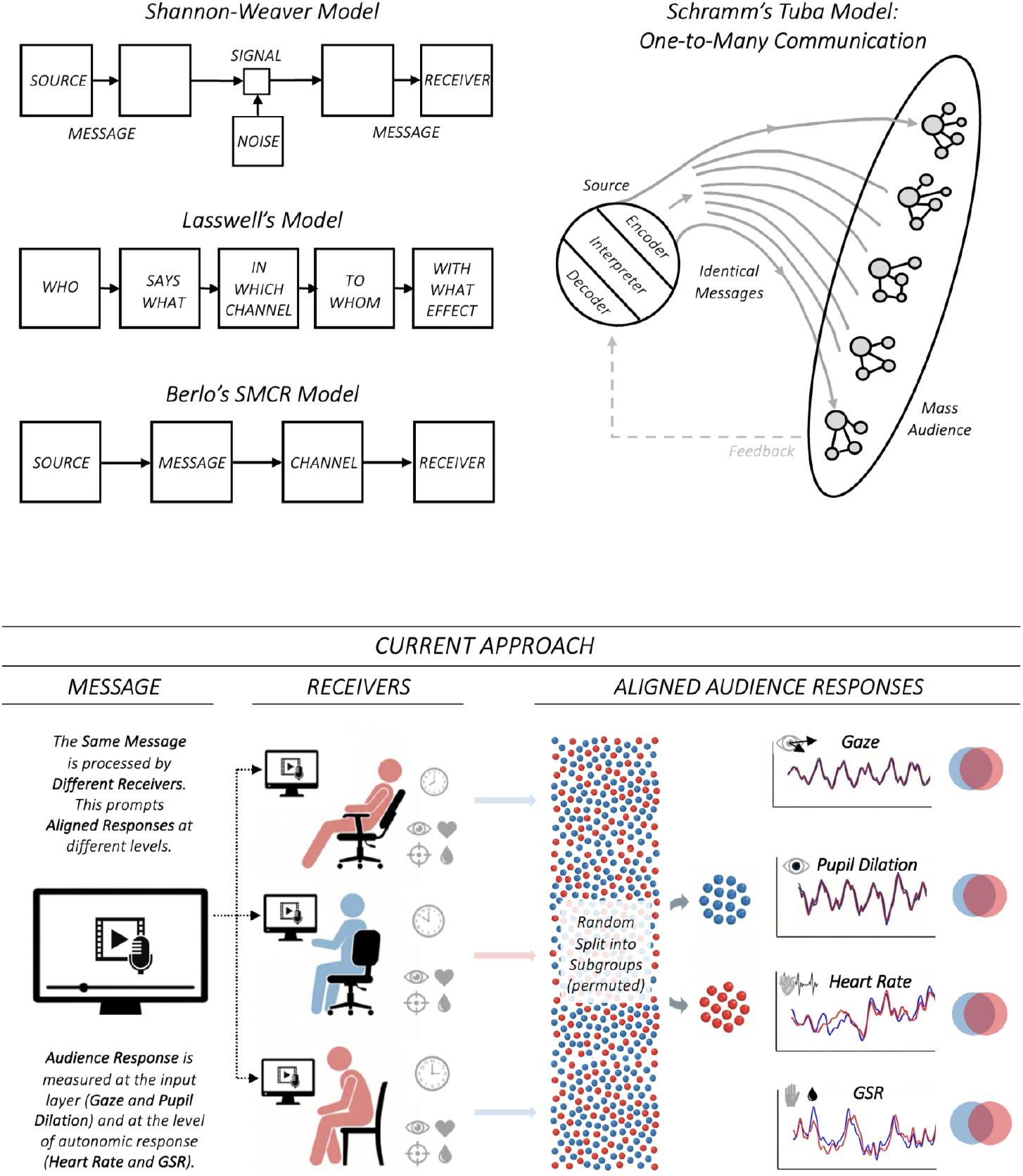
Historical Communication Models and Illustration of Audience Response Alignment. Early models (e.g., Berlo’s SMCR) emphasized the transmission of information from source to receiver. Schramm’s Tuba Model specifically highlights the “one-to-many” dispersal of a signal to a multiplicity of receivers. The current framework investigates the fidelity of this transmission by measuring physiological alignment. Regardless of whether viewing is asynchronous or remote, the narrative signal entrains the audience’s biological rhythms, from information acquisition (gaze direction, pupil dilation) to autonomic regulation (heart rate, GSR). These convergent responses demonstrate that the audience is not merely a statistical aggregate, but a functionally defined group of receivers brought into temporary inter-subjective alignment by the temporal structure of the message.

### From Neural Processing to Bodily Resonance

The mechanism supporting this alignment is the shared biological architecture of the human receiver. As posited by the Extended Neurocognitive Network Model (Schmälzle, 2022), humans possess functionally homologous sensory and processing hardware. This shared architecture implies that a standardized input signal, if sufficiently engaging, should elicit converging audience responses across individuals who process the same message. While neuroimaging research has firmly established that messages drive reliable alignment in the central nervous system (brain synchrony; Hasson et al. (2004); Nastase et al. (2019)), we argue that the “physiological reach” of a narrative message signal extends beyond the brain (cf. Green et al. (2002)). If a message effectively recruits the cognitive and emotional machinery of the receiver, this alignment should propagate downstream to the peripheral nervous system. By measuring these peripheral outputs, specifically the viscero-motor regulation of the heart and skin, we can trace the depth of the signal’s reach, asking whether the message is powerful enough to entrain not just the neural processor, but the homeostatic rhythm of the body itself.

### Hierarchy of Audience Alignment: From Information Acquisition to Autonomic Regulation

While the shared biological architecture suggests a capacity for alignment, the manifestation of this synchrony is not uniform. The human system operates hierarchically (Mesulam, 1998), implying that a message must traverse a processing cascade: it must first capture sensory inputs before it can drive higher-order cognitive^3^, emotional, and autonomic states. Consequently, we posit that audience alignment manifests in two distinct layers: Attentional alignment (information acquisition) and affective resonance (autonomic regulation).

The first layer, attentional alignment, is visible in the synchronization of gaze behavior. In audiovisual narratives, eye movements are heavily constrained by formal features such as editing, motion, and luminance, a phenomenon described as the “tyranny of the film” (Loschky et al., 2015). High gaze alignment serves as the necessary gateway for communication,confirming that the filmmaker has successfully standardized the perceptual input channel across the group (Madsen et al., 2021; Madsen and Parra, 2022).

The second, deeper layer is affective resonance, measured via autonomic outputs (heart rate, skin conductance; Golland et al. (2014); Han et al. (2022)). Unlike gaze, which is largely driven by bottom-up sensory features, autonomic responses are continuously modulated by internal homeostatic processes, individual states (e.g., fatigue, arousal baseline), and psychophysiological modulations Ravaja (2004); Potter and Bolls (2012); Berntson et al. (2017)). Therefore, achieving audience alignment at this level represents a more demanding test of signal fidelity. If a story can elicit time-locked fluctuations in heart rate (HR) or skin conductance responding (SCR) across a group, it demonstrates that the message signal is robust enough to overcome the receiver’s individual variance and entrain viscero-motor systems that are key components of emotional responses (Lang, 1988).

In summary, this hierarchical framework leads to a specific prediction regarding the magnitude of audience alignment. We expect a gradient with extremely high correlations for information acquisition (Gaze/Pupil), reflecting the “tyranny” of the visual input, followed by lower but significant correlations for autonomic outputs (HR/GSR), reflecting the successful induction of a shared affective states despite individual biological variability.

### Signal Specificity of Message-Induced Audience Alignment

Demonstrating that audiences exhibit convergent physiological responses raises a critical question regarding the nature of this alignment: Is it a lingering, blurry, and tonic state dependent on the ongoing mood or global atmosphere of the story, or is it a robust, time-locked and more phasic response to specific audiovisual inputs? Arguably, to serve as a more interesting measure for communication, the physiological signal should exhibit temporal specificity. That is, the response should not merely be similar across viewers in a general sense, but it should be directly linked to the specific moment-to-moment unfolding of the content. This distinction is crucial for ruling out alternative explanations. If the physiological convergence observed in an audience were merely the result of a generalized emotional atmosphere (e.g., the audience simply feeling excited throughout the film), the response would lack temporal precision. In that case, swapping a segment might change the overall level of arousal, but it would not produce a distinct, time-locked pattern that tracks with the swapped content. However, if the response is driven by a robust coupling between the stimulus and the receiver, a sort of physiological (temporal) fingerprint of the content itself, then the alignment should prove resilient. Specifically, the physiological signature of a scene should “travel” with that scene, regardless of its position in the timeline. If the physiological response can provide such a physiological fingerprint or signature of a scene, then this characteristic response should persist even when that scene is moved to a different point in the story timeline. To empirically isolate this robustness and temporal (scene) specificity, it is necessary to decouple the content from its chronological position:

If the alignment is truly content-locked, we would expect two distinct result patterns under manipulation: First, when two audiences view different message segments at the same absolute time, their physiological alignment should vanish (demonstrating that the signal is not random). Second, when the physiological time-series are computationally realigned to match the content (effectively “unscrambling” the response based into the original order of the scene sequence, Hasson et al. (2008b)), then the correlation between the groups should be restored. Validating this robustness would demonstrate that the convergent response is a reliable, replicable signature of the message that persists even under disrupted conditions. Ultimately, this would confirm the extensive biological impact of narrative messages: demonstrating that a single audiovisual signal can propagate through the idiosyncrasies of distinct viewers, regulating their physiology with enough precision to align their responses - from gaze patterns to even their autonomic outputs like heart rhythms and sweat gland activity - into a shared physiological state.

### The Present Study and Hypotheses

To empirically test the reach and robustness of this audience alignment, we conducted a large-scale experiment (N = 198) using continuous psychophysiological tracking. Participants viewed Pixar animated short film, Partly Cloudy (2009) while their information acquisition (Eye Movements, Pupil Size) and autonomic regulation (Heart Rate, Electrodermal Activity) were recorded.

Inspired by previous work using scrambling to alter temporal structure (Lerner et al., 2011), we employed the following message manipulation: First, we analyzed a main audience group exposed to the unaltered story to establish the presence and hierarchy of convergent audience responses (Condition: PC). Second, we compared these responses to those from a group of viewers exposed to a modified version of the story in which the climax and resolution segments were swapped (Condition: PCM).

Based on the theoretical framework outlined above, we posit the following hypotheses:

#### H1 (Hierarchy of Alignment

Message exposure will induce an aligned (synchronized) physiological state across the audience, manifesting as significant Inter-Subject Correlation (ISC) across all measures. We predict a hierarchical gradient, such that alignment will be strongest for information acquisition channels (Gaze/Pupil) and lower, yet significant, for autonomic output channels (HR, GSR). This hierarchy reflects the attenuation of the external signal as it moves from standardized sensory input to the internal regulation of the viscero-motor system.

#### H2 (Temporal Specificity)

Audience alignment is dependent on the temporal synchronization of content. Therefore, during message segments where the two groups view different content (i.e., the swapped climax and resolution), ISC calculated between the main and manipulated groups will be significantly reduced or abolished compared to segments where the content is identical.

#### H3 (Signal Robustness)

The physiological response to the message serves as a content-locked temporal fingerprint of the message. Therefore, computationally unscrambling the time-series of the group that saw the edited/modified movie (i.e., realigning their physiological data to match the chronological order of the main group) will restore significant inter-group correlation.

#### RQ1 (Dynamics of Divergence)

While H2 predicts that divergence will occur, it does not specify how the biological system responds to the moment of disruption. Given that autonomic processes (e.g., sympathetic arousal) often exhibit slower decay rates than attentional processes (e.g., saccades), we ask: How quickly does physiological alignment dissolve following a temporal structure manipulation, and does this rate of divergence differ between attentional (Gaze) and autonomic (HR, GSR) measures?

## Methods

### Participants

A total of 208 participants were initially recruited for the study. Ten individuals were excluded and replaced prior to the experimental task due to failed eye-tracking calibration (*n* = 6), or technical equipment failures (*n* = 4; e.g., disconnection of the headset, camera, or computer), resulting in a final sample of 198 participants (*mean*_*age*_ = 19.8, *sd* =1.7, 105 self-identified males). Half of the sample (99 each) viewed either the normal, unedited version of Partly Cloudy (5 min 19 sec version, *PC*) or a manipulated version in which the climax and resolution segments were switched (details see below, *PCM*). All participants provided written informed consent to the study protocol, which was approved by the local review board. They received course credit for their participation in the study, which took about 45 minutes.

### Stimulus

#### Partly Cloudy (PC) - Original Version

The study utilized the dialogue-free Pixar animated short film, Partly Cloudy (2009) as a stimulus. This short film has been previously used to study theory-of-mind and emotional processing (Richardson et al., 2018). The plot follows Gus, a cloud, and Peck, his stork delivery partner. Conflict arises because Gus creates hazardous “babies” (e.g., sharks, porcupines) that injure Peck. Peck’s eventual flight to another cloud hurts Gus emotionally, making him cry. The film concludes with Peck’s return and a positive resolution of their conflict, highlighting themes of loyalty, friendship, and mutual understanding.

#### Manipulated Version (PCM)

To create a temporally manipulated version of this movie, we followed research that analyzed this movie and broke it down into story segments (premise, rising action, climax, and resolution) (Grady et al., 2022). We then kept the initial two segments in the same order but switched the order of the resolution and the climax. This created a movie that was still continuous and comprehensible, presenting the same characters and actions throughout, although it switched from an ending in which Peck returns to Gus and they reunite to one where the conflict goes on and Peck eventually leaves.

### Procedures

Participants watched the movie on a 24-inch monitor placed about 60 centimeters in front of them. Before the movie started, they filled out consent and were prepared for the physiological measures (see below). To ensure data quality for eye-tracking, an eye-tracking calibration was performed using iMotions software; participants were required to achieve a calibration quality of “good” or “excellent” before proceeding. Those who failed to meet this calibration threshold (*n* = 6) were excluded and replaced, as noted in the Participants section. Following successful calibration, they watched a 1.5-minute video before the Partly Cloudy movie started. The movie was shown without specific instructions (free viewing). After the movie ended, participants filled out brief questions regarding their prior familiarity with the stimulus, comprehension, attention, entertainment experiences, emotional responses, as well as narrative engagement.

### Measures and Analysis

#### Data Collection

Physiological data were recorded on two iMotions workstations equipped with Smart Eye Aurora for screen-based eye tracking, Shimmer 3 GSR kit and Shimmer 3 ECG kit. Data were sampled at 60 Hz for eye-tracking, 128 Hz for GSR, and 512 Hz for ECG.

We also collected post-hoc self-report data after participants had watched the movie. This included a single-item measure of prior exposure (i.e., whether they had seen the film before), the emotional flow scale (Fitzgerald et al., 2023), summary-style measures of enjoyment and appreciation (adapted from Oliver and Bartsch (2010)), attention, and narrative engagement (Busselle and Bilandzic, 2009).

#### Data Export, Preprocessing, and Analysis

All analyses were carried out in Python and are documented at [blinded for review]. Data were cleaned, preprocessed, and exported using iMotions’ validated processing routines. We began the analysis by reading in each subject’s data for each response channel and clipping out the parts before/after the movie (based on iMotions triggers). Specifically, for eye-tracking, we read in the x-and y-Gaze coordinates and the pupil diameter, both for the left and right eye, which were averaged. Heart Rate (HR) was converted to Beats Per Minute (BPM) and resampled to 2 Hz; Electrodermal activity was high-pass filtered with a 0.05 Hz Butterworth filter to focus on phasic components. Slow signals (HR and GSR data). Given that HR operates within a physiologically constrained range (typically 60-120 BPM), raw BPM values were preserved to maintain biological interpretability and only normalized for plotting purposes.

To prepare the main dataset, data were then organized by subjects, by condition, and by response channel, resampled to a common 20 Hz sampling and saved as a single file with a common timeline.

Next, we examined the timeseries and computed inter-subject correlation analysis following prior work (Hasson et al., 2004; Nastase et al., 2019; Golland et al., 2014). First, to address H1, we computed separate ISCs for each response channel (Gaze, Pupil, HR, GSR) for the entire film. We examined the data using a 5000-fold permuted split-half analysis for signal-plotting and inspection, followed by a pairwise ISC analysis with a 5000-fold phase-randomization procedure to demonstrate that the observed ISC is robust when compared against surrogate data whose temporal alignment has been destroyed (see Nastase et al. (2019)).

Next, to address H2 and H3, we further separated the physiological time series based on the movie segments and computed ISC analyses both within and across the two viewing groups exposed to the original or the modified movie version. Thus, we computed ISC for those segments that were identical across the two movie viewing conditions (i.e., beginning, premise, rising action, ending) and also for the segments where the temporal/chronological order had been reordered (i.e., climax, resolution, which were switched for the reordered movie), assuming that the latter manipulation would cause the ISC between groups (original/main movie audience vs. edited/manipulated movie audience) to drop. Additionally, we also computed a dynamic ISC analysis in which we used a sliding window procedure to examine whether dynamic ISC would evolve similarly with the shared underlying message and then start to diverge as the message diverges.

To test the fingerprint hypothesis (H3), we performed a temporal realignment procedure: The physiological time-series data for the manipulated group were computationally cut and re-ordered to match the chronological sequence of the original/main group. This allows us to test if the physiological response pattern travels with the story content, independent of its position in the viewing timeline.

Lastly, to address the research question, we ran cross-correlations between the two groups using a forward (causal) sliding window ISC procedure in order to test how quickly the convergent audience responses diverge and fall apart as the story branches.

Above and beyond these main analyses, we conducted additional control and extension analyses that are reported in the Supplement. These include specifically a reverse correlation analysis that convincingly demonstrated that salient events in the story drove collective peaks in the GSR response across the audience, that prior knowledge of the story did not confound results, that attentional alignment between individuals predicted autonomic alignment, and that dynamic ISC between the main and manipulated audience evolved similarly (further supporting H2/H3).

## Results

Analysis of post-viewing surveys confirmed that participants in both conditions successfully comprehended the plot and reported high levels of emotional flow (*m*_*PC*_ = 4.97, *m*_*PCM*_ = 4.76), as well as ratings of enjoyment (*m*_*PC*_ = 5.83, *m*_*PCM*_ = 5.4) and appreciation (*m*_*PC*_ = 4.54, *m*_*PCM*_ = 4.3). Participants reported taking interest in the movie and paying close attention overall (*m*_*PC*_ = 5.94, *m*_*PCM*_ = 5.85) and being engaged (*m*_*PC*_ = 4.99, *m*_*PCM*_ = 4.65; max = 7). With valid reception established, we now turn our focus to the real-time dynamics of the audience response. The upcoming results are presented in two streams:

The first stream of analyses characterizes the collective physiological alignment of the audience and establishes the hypothesized hierarchy of physiological alignment (Gaze > Pupil > HR/GSR) within the group exposed to the main movie stimulus (PC). Then, a follow-up analysis stream delves into the temporal nature of this alignment, comparing the baseline and manipulated groups to test the robustness and content-specificity of the physiological signal.

### Alignment of Main Audience

As expected, and shown in Figure 2, the movie prompted not only highly similar gaze patterns, but it also consistently modulated Pupil Size, Heart Rate, and GSR. Figure 2 shows split-half plots that were created by randomly splitting the audience (97 viewers after dropping 2 because of a bad signal in one of the 5 measures) into two groups and averaging their data. Of note, because there are many ways to split 97 viewers into two groups, the correlations shown in Figure 2 are subject to noise. To overcome this, we ran 5000 permutations and averaged the results, which confirmed the one split illustrated in Figure 2 and yielded the averaged ISC-values mentioned next: We find that the Gaze coordinate time courses (X and Y signals), exhibit extremely high correlations of _*rGazeX:Split-Half ISC*_ =.96 (CI: [.953,.974] and *r*_*GazeY:Split-Half ISC*_=.95 (CI: [.936,.959]). Similarly, the pupil size (*r*_*PupilSize:Split-Half ISC*_ =.962, CI: [.942,.972]), the heart rate (*r*_*HeartRate:Split-Half ISC*_ =.702, CI: [.554,.797]) and the skin conductance (*r*_*GSR:Split-Half ISC*_ =.514, CI: [.401,.609]) all are robustly correlated across the audience. This pattern of responses also confirms the predicted response hierarchy: Gaze and pupil-measures, which are close to the stimulus input, are most highly correlated, followed by heart rate and the skin conductance, which represent more downstream responses.

**Figure 2.**
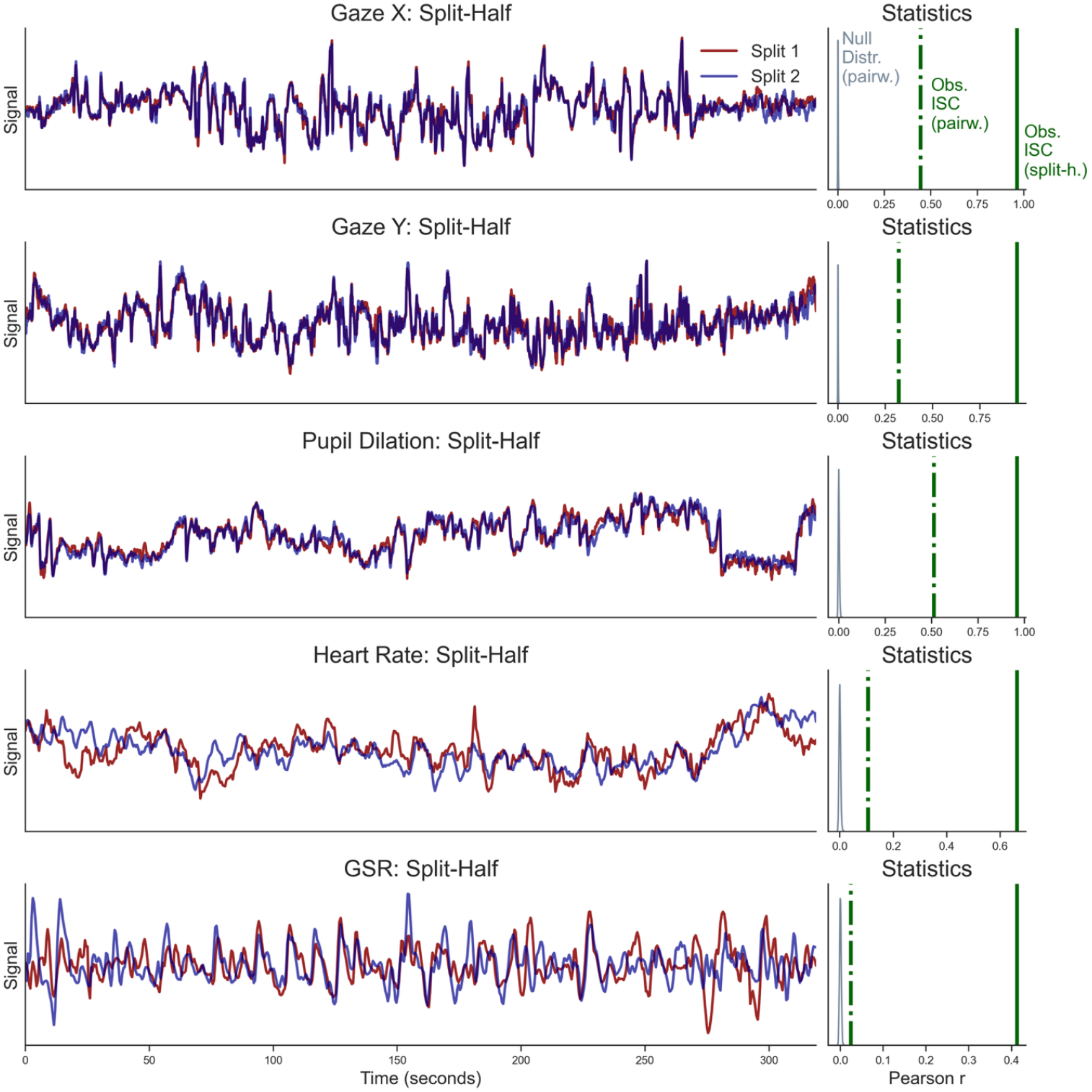
Alignment of Audience Responses for Main Audience. Plots on the left side show averaged time course for all channels based on a split-half analysis in which the audience is randomly split into two groups whose responses are then averaged. The plots on the right side illustrate the actual, observed ISC (green lines, dashdotted: pairwise and solid: split-half ISC metrics) as well as null-distribution obtained via a 5000-times phase-randomization procedure. As can be seen, ISC is highly significant across all response channels but varies in magnitude from very high (gaze, pupil, >.9 for split-half comparison), to medium (ca..7 for heart rate), to moderate (ca.4 for GSR).

Beyond these split half analyses, we also computed statistical tests to ensure that the synchronization observed was robust and different from zero. Using the pairwise ISC with a 5000-fold phase-randomization analysis fully confirmed the pattern seen with the split-half method used for visualization: The ISC was highly significantly positive for all five measures (Gaze X and Y trajectories, *p*_*gazeX*_ <.0002; *r*_*GazeX:pairwise*_ =.42; *p*_*gazeY*_ <.0002, *r*_*GazeY:pairwise*_= 0.31; Pupil Size *p*_*PupilSize*_<.0002, *r*_*PupilSize:pairwise*_=.5;Heart Rate *p*_*HeartRate*_ <.0002, *r*_*HeartRate:pairwise*_ =.11; and GSR *p*_*GSR*_ <.0002, *r*_*GSR:pairwise*_ =.026) and destroying the temporal alignment of the physiological timeseries (via 5000 fold randomization statistics) made ISC vanish. In sum, these results support H1.

### Temporal Specificity of ISC

Next, to test H2 that focused on the temporal specificity of ISC, we computed ISC not only for the main movie and its audience, but also for the modified version, taking into account the segments that were either identical across both movie versions/audiences (i.e. the first half of the movie, containing the premise and rising action phase, was identical in both versions, and also the ending credits) or different (i.e. while the main movie proceeded normally, the modified version had a scene switched, which should break the temporal alignment). Thus, we computed the same analyses as done above for the main movie, but now for the subsegments that were common or not common between the two audiences. As shown in Figure 3, the common segments prompted responses that were highly correlated across the two sub-audiences and similar in magnitude as for the main analysis reported above(*r*_*GazeX:Split-Half ISC Across Group*_= 0.96; *r*_*GazeY:Split-Half ISC Across Group*_=.95, *r*_*PupilSize:Split-Half ISC Across Group*_=.95; *r*_*HeartRate:Split-Half ISC Across Group*_=.76, *r*_*GSR:Split-Half ISC Across Group*_=.4). Note that this analysis again replicates the gradient of ISC strength, ranging from very high gaze-and pupil ISC (above.9)to lower but still robust ISC for HR and GSR (between.4-8).

**Figure 3.**
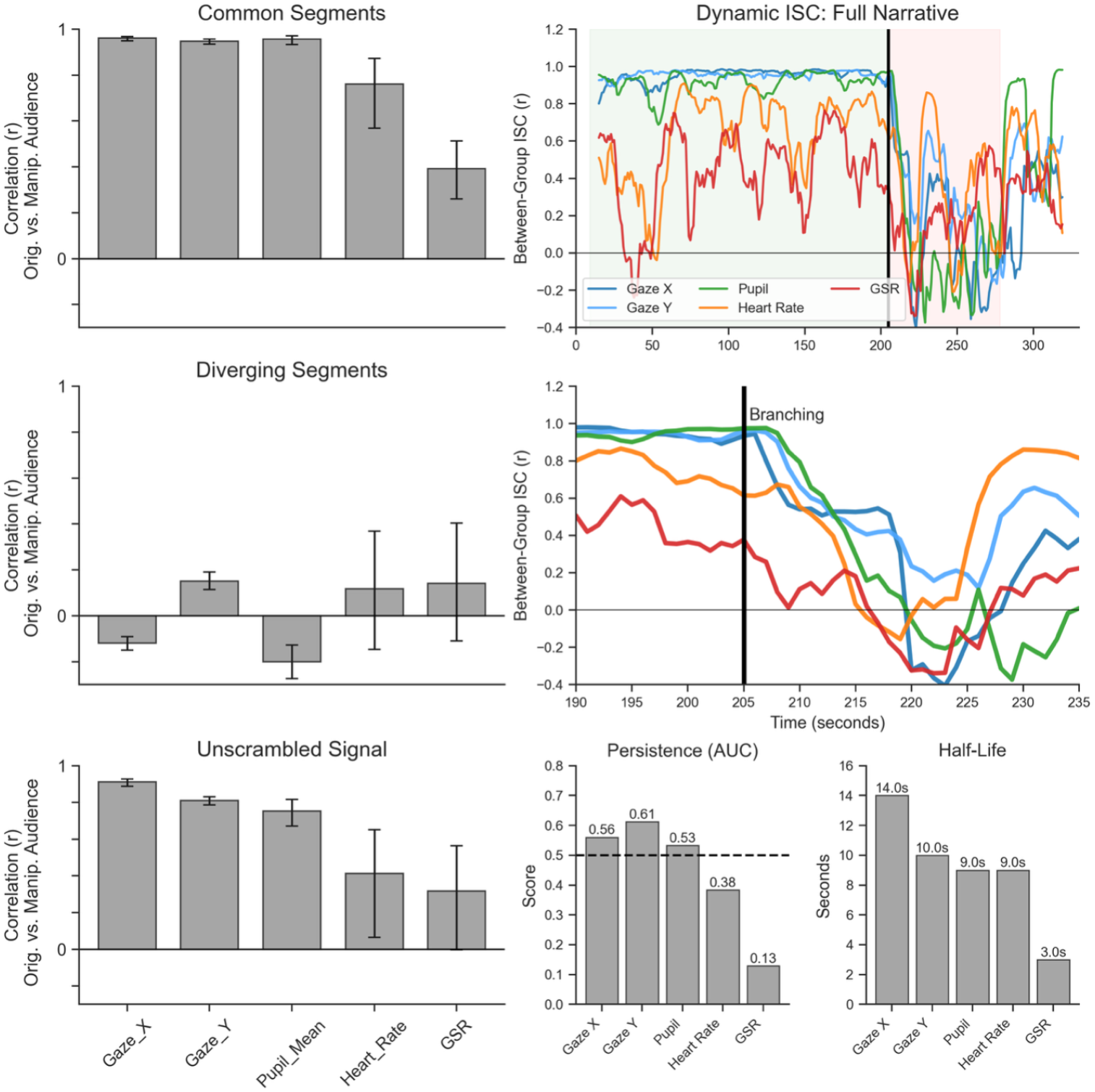
Convergence and Divergence of Audience Responses: Left: Shared responses for common segments (top), alignment break during story branching (middle), and computational realignment (bottom). The graphs and variability metrics are based on a 500-times permuted split-half ISC analysis. Right: Sliding-window ISC tracks the dynamic evolution of ISC across the shared segments and documents its rapid decay after the story branching. Top right: A sliding window ISC is computed between the data from one audience and the data from the second audience.

Moving on to the critical branching of plots, we find a stark contrast: The entire pattern of common responses breaks down when the scenes are not aligned, and some of the raw ISC values are even negative (see Figure 3, middle panel). This demonstrates that the ISC between the two sub-audience is driven by the common input: if the input is the same, we see common responses among all people exposed to the original movie, among those exposed to the edited version (which in the beginning is identical to the original, however), and also across those groups. However, if the input diverges, audience responses diverge. These results support H2.

Lastly, we turn to H3, which stated that computationally unscrambling the time-series of the group that saw the manipulated movie sequence (i.e., realigning their physiological data to match the chronological order of the main group) would restore significant inter-group correlation. The results of this investigation are shown in the lower panel in Figure 3. As can be seen, computational unscrambling restored the alignment of physiological responses across group and to almost the same level, with gaze ISC going back to *r*_*GazeX:Split-Half ISC Across Group*_ =.91 and *r*_*GazeY:Split-Half ISC Across Group*_=.81, respectively and ISC values of *r*_*PupilSize:Split-Half ISC Across Group*_ =.7, *r*_*HeartRate:Split-Half ISC Across Group*_=.4, and *r*_*GSR:Split-Half ISC Across Group*_= 0.3 for the pupil, heart rate, and GSR, respectively. These results support H3.

### Dynamics of Divergence Between Main and Manipulated Audience

Finally, to address our research question, we explored the temporal dynamics of desynchronization following the narrative disruption (Figure 3, right panel and Figure S1). Inspired by Excitation-Transfer Theory (ETT), which suggests residual arousal lingers across events, we used a causal (i.e., forward-looking window) sliding-window ISC analysis to track how quickly the shared signal dissolves once the storylines diverge. We found that measures of gaze synchronization persisted for several seconds (consistent with the mechanical delay of the sliding window and the movie’s continuous flow), but physiological alignment evaporated more quickly (although it should be noted that this also started from a lower level and generally exhibited more fluctuations over the movie). GSR, in particular, exhibited a half-life of only 3 seconds and a Persistence Score of just.10 (where.50 represents a neutral linear decay). Thus, contrary to the hysteresis (or memory, lag) predicted by Excitation-Transfer accounts, this suggests an active overwriting effect: the incoming divergent content immediately commanded specific physiological regulation, effectively erasing the shared arousal state rather swiftly as the scenes changed. It should be noted, however, that ETT focuses usually on salient moments, whose excitation then carries over to more standard content. The case studied in this exploratory analysis, however, refers to a more normal scene transition (albeit at the rising action phase of the movie) that is followed by a normal and continuous narrative that fits the storyline in both versions.

## Discussion

In this investigation, we examined the extent to which a narrative signal can coordinate the biological states of distinct individuals to induce a convergent audience response. Using a large-scale experiment with continuous psychophysiological tracking, we found that the message successfully entrained not only the visual attention of the audience but also their downstream autonomic regulation. These findings expand the traditional view of the audience as a mere statistical aggregate of demographic clusters. Instead, our data support a bio-functional definition: the audience is a temporary network of receivers brought into existence and functionally aligned by the temporal structure of a message.

### Key Results

#### Aligned Responses

Our first finding establishes the robustness of convergent audience responses as well as their proposed hierarchical nature (H1). As predicted, we observed a gradient of synchronization: synchronization was highest for information acquisition behaviors (Gaze/Pupil, *r* >.90) and lower, yet robustly significant, for autonomic outputs (Heart Rate, *r* ca..70; GSR, *r* ca..50; split-half method).

#### Hierarchy of Signal Propagation

This hierarchy maps the processing cascade of communication. The extremely high gaze alignment confirms the tyranny of the film (Loschky et al., 2015), demonstrating that formal features (motion, editing) successfully standardized the perceptual input channel across the group. However, the significant alignment of downstream autonomic measures represents a more profound communicative feat. Heart rate and skin conductance are typically regulated by internal homeostatic needs (e.g., thermoregulation, metabolic state) as well via complex psychophysiological processes (Potter and Bolls, 2012; Ravaja, 2004). The fact that an external audiovisual signal could couple these systems across distinct individuals suggests that the message signal was strong enough to overcome the biological idiosyncrasies of receiver. While gaze alignment indicates people in the audience looked at the same things, autonomic alignment implies they experienced a shared fluctuation of arousal, effectively reducing the variance between distinct mind/body systems and promoting shared reality (Echterhoff et al., 2009).

#### Signal specificity

A common critique of psychophysiological measures in media research is that they may simply reflect a generalized atmosphere or tonic state of arousal induced by the genre. Our results refute this interpretation through the demonstration of temporal specificity (H2) and signal robustness (H3). When the message structure was broken (by swapping the climax and resolution), the physiological alignment between groups vanished (H2). This divergence confirms that the observed synchrony is linked to the specific moment-to-moment unfolding of the content. Most compellingly, computational unscrambling of the data restored the correlation (H3, almost as high as the original values). This finding supports the physiological fingerprint hypothesis: specific narrative events (e.g., a moment of conflict or resolution) elicit reliable, phasic physiological signatures that “travel” with the content. This implies that the biological response is content-locked, it is a reaction to the information provided by the signal, not a random fluctuation of the receiver’s mood.

### Implications for Communication Theory and Media Practice

These findings have broad implications for how we conceptualize and examine the audience in communication research. For decades, the field has moved away from transmission models to focus on the active, interpretive role of the receiver. While this shift is necessary to understand socio-cultural phenomena, it has perhaps led to an underestimation of the signal’s organizing power in the first place. Our data suggest that the audience can not only be understood as a sociological category, but can be productively studied as a functional state of inter-subjective alignment that is actively induced and maintained by the message.

Our findings also provide empirical support for recent theoretical work conceptualizing communication as a mechanism of alignment. Gasiorek and Aune (2021) propose that successful communication results in the alignment of minds, a state where the cognitive and semantic representations of interlocutors become synchronized. Our results suggest that this alignment is not limited to high-level cognition but reaches deep into the regulation of the nervous system, literally down to the fingertips. Before minds can align semantically, bodies must align physiologically.

Theoretically, this relates to the Shannon-Weaver model Shannon (1948), anchoring it at a biological level. We acknowledge that applying information theory to human communication requires caution. In his editorial *“The Bandwagon,”* Shannon (1956) explicitly warned against the loose application of engineering concepts to the social sciences, fearing that the mathematical precision of the theory would be diluted when applied to complex, subjective human behaviors (see e.g., Schramm (1955) article *“Information theory and mass communication”*). For decades, this warning remained prescient, as tension and theoretical or even paradigmatic commensurability gaps continue to persist between communication engineering and communication studies. However, the current biofunctional approach offers a potential way forward that many theorists would likely support. By focusing on physiological responses rather than interpretations, we move the level of analysis from a subjective level to an objective one dealing with the accuracy of transmission.

In this context, ISC serves as a mathematical quantification of fidelity. When we observe high inter-subject correlations, we demonstrate that the specific temporal structure of the message has successfully propagated through the biological channel of the receiver. This reinforces the utility of signal-processing models in communication science (Cherry, 1966). It suggests that a successful message is one that possesses sufficient signal fidelity to penetrate the “noise” of individual difference, establishing the material foundation for communication to function in society. Theoretically, it relates to the concept of shared reality (Echterhoff et al., 2009), but with an emphasis on material substrate (Hasson and Frith, 2016). Before people can share a subjective understanding of a message, they must share a processing state. The high-fidelity propagation of the message signal, from visual input down to autonomic regulation, provides this necessary condition. It reduces the biological uncertainty between distinct individuals, creating a shared state that forms the basis of higher-order meaning construction and collective experiences. This perspective invites a re-evaluation of the “Signal-to-Noise” ratio in human communication. In this context, noise is not merely channel interference, but the biological and psychological variability of the receiver (e.g., individual differences in attention, mood, or background; Nastase et al. (2019)). A successful narrative acts as a high-fidelity signal capable of overcoming this receiver-side noise. The fact that we observed such physiological alignment across a diverse sample (*N*=198) suggests that when the signal is sufficiently structured, the specific attributes of to whom the message is sent become less predictive of the processing state than the message itself. This does by no means negate the importance of individual differences or notions like audience activity; rather, it highlights the impressive efficacy of narrative form in coordinating the human organism to make mass communication possible.

To avoid misunderstanding about these issues, two remarks are in place: First, readers may wonder to what extent these findings, which were obtained with a narrative film that is presumably rather optimized for audience engagement generalize to other kinds of mass media messages? This is indeed a valid point and one that can and should be studied empirically (e.g., Hasson et al. (2008a) on the study of movies, Schmälzle et al. (2013) for a study of TV documentaries, Grall et al. (2021) for radio stories; Jangraw et al. (2023) for written messages). Thus, while the argument that the current work relies on a single stimulus that may not represent the entire genre nor the entire universe of messages, the more general point about the nature of convergent audience responses during the reception of the same mass communication messages still stands. The second point is about the complementary nature between commonalities and differences. As stated above, the finding that there are convergent audience responses, which evidently matter for mass communication, does not negate individual differences. Indeed, prior work has already shown differences in message reception (e.g. pre-existing knowledge, attitudes, traits) can be studied using the same theoretical toolkit, basically enabling physiological audience segmentations. Thus, the focus here is on the commonality in inter-subjective message reception processes and physiological effects, but with larger samples and adequate messages one can also study divergent audience responses.

### Practical Implications: Signal Fidelity in a Fragmented Media Landscape

Practically, the robustness of the physiological fingerprint (H3) offers critical insights for the modern media landscape. In an era of fragmented consumption, where narratives are often consumed in non-linear clips via social feeds (e.g., TikTok, YouTube Shorts), on different devices and in different settings, the question of whether context matters is paramount. Our results suggest that the physiological signature of engaging content remains intact even when decoupled from the global narrative arc. If a segment possesses high signal fidelity, it creates a convergent response even in the absence of the broader story. This validates the efficacy of short-form storytelling: it suggests that high-density narrative signals can create “micro-audiences”, groups functionally aligned by the content, even in snippets (cf. Schmälzle et al. (2025)). The potential to measure these micro-level alignments via wearables (as demonstrated by our use of standard sensors) opens new avenues for understanding audience responses and engagement in real-time.

Finally, this robustness also implies that we can treat measured audience responses as a continuous annotation of the content. In the field of deep learning, the availability of labeled data has been the driver of major advances (Sambasivan et al., 2021), be it in the form of language annotations or reinforcement learning from human feedback. With this in mind, we see a bright future for media psychophysiology because it allows us to capture how audiences respond to movies and link responses back to their elicitors in content (Schmälzle et al., 2022) or to predict out-of-sample responses (e.g., (Dmochowski et al., 2014; Falk et al., 2015)), which has many applications beyond the theoretical insights foregrounded here.

### Limitations and Conclusion

This study is not without limitations. First, while we quantified the synchrony of the processing state, we did not measure the semantic content of the audience’s subjective experience. Future research should bridge this gap by combining continuous physiological tracking with continuous semantic annotation (e.g., perception analyzer dials, Levy (1982)) to map how biological entrainment supports specific interpretations. Second, our stimulus was a dialogue-free animated film; while this maximized cross-cultural validity and focuses on the long-known strong effect of moving images and minimizes linguistic decoding noise (Münsterberg, 1916; Bente et al., 2022; Baldwin and Bente, 2021), future work should examine how verbal content interacts with this physiological layer. Third, we used only one stimulus film and only made one scene-swap. While these choices were justified given the nature of the research, future research should examine multiple stimuli and conduct additional tests of scrambling and the signature/fingerprint analyses, especially regarding the connection to Excitation-Transfer theory. Given that narratives are inherently temporal in nature, manipulations like the ones we suggest would seem ideally suited to test hypotheses about emotional flow and related dynamic emotion phenomena. Another avenue that seems promising would be to give different participants different tasks (e.g. attentive watching, watching with a particular perspective or pre-knowledge, or group-membership; Ki et al. (2016); Branford and Johnson (1972); Hastorf and Cantril (1954); Lazarus et al. (1962) and examine the effect on convergent audience responses.

In conclusion, we show that the audience is not merely a statistical abstraction or a demographic cluster. When we observe high correlations in attention and autonomic regulation measures, we can prove that the message signal has successfully crossed the gap between isolated receivers. By measuring the propagation of the narrative signal through the viewers’ hierarchy of processing, we demonstrate that mass communication acts as a powerful organizing force on the receiver system (individually and collectively). A successful message aligns the audience as a group into a shared functional state. In doing so, it creates the material foundation for shared reality, bridging the gap between isolated individuals.

## Declarations

### Ethics approval

The study was approved with the IRB of [blinded for review].

### Consent to participate

All participants provided written informed consent.

### Use of Generative AI

Google Gemini and Nano Banana were used during the drafting process to assist with the organization of the manuscript, the refinement of theoretical arguments, copy-editing, and creating elements of the figures. All original ideas, hypotheses, data analysis, and final theoretical conclusions remain in the intellectual property of the authors. The authors reviewed and edited all text and take full responsibility for the content.

### Availability of data and materials

All data and materials, except for private raw recordings (video/audio), are available at [blinded_review].

## Supplementary Materials

### S1 Reverse Correlation of GSR Responses

To investigate the observed physiological alignment and identify the specific narrative drivers of autonomic arousal, we employed a reverse-correlation approach (e.g., Hasson et al., 2004; Schmälzle et al., 2022). This analysis aimed to work backwards from the biological response to the narrative stimulus, verifying that moments of shared GSR peaks were not random but driven by salient narrative events.

First, we isolated the significant peaks in the group-averaged GSR signal. We calculated the group average across all participants in the main condition to maximize the signal-to-noise ratio and isolate the shared component of the audience response. Peaks were identified using the find_peaks algorithm (SciPy signal processing library) with a minimum prominence threshold to filter out minor fluctuations.

Second, to account for the known physiological latency of the electrodermal response, we applied a temporal shift operation. As GSR peaks typically occur 3-6 seconds following a stimulus (Boucsein, 2012), we defined the event time t(event) as the physiological peak time *t(peak)* minus a fixed lag of 4.5 seconds. We then extracted 3-second video clips centered on these trigger times, i.e., *t(trigger)* = +/– 1.5s. We also generated a matched set of control clips, which were extracted from random timestamps within the movie, with the constraint that they could not overlap with the identified peak windows.

We then inspected the extracted segments (*n* = 8) visually and found that the peak-clips contained two highly salient scenes one would describe as definitely high-arousing: The first scene is where Peck, the stork, gets his head almost bitten off by a baby crocodile (see Figure S1), the second scene identified from the GSR peak is when a baby ram crashes into Peck. Above and beyond this visual inspection, we also attempted to further quantify this by utilizing a Large Multimodal Model (Gemini 3 Pro) as a blind coder. The model processed both the peak (*n* = 8) and control (*n* = 8) clips in a randomized order and rated them on a scale of 0-100 for Narrative Intensity, Physical Threat, and Emotional Arousal. We then created a summary score and compared scores for peak and control clips via an independent sample t-test. The average score for peaks was 49.8 (*sd* = 27.6), the average for controls was 30.2 (*sd* = 18), which is not a significant difference (*p* = 0.058), although the low sample size must be considered as a hard hurdle to overcome. Overall, on top of our own visual inspection, this computational approach provides an independent validation that the physiological peaks were driven by objectively intense narrative moments.

**Figure S1.**
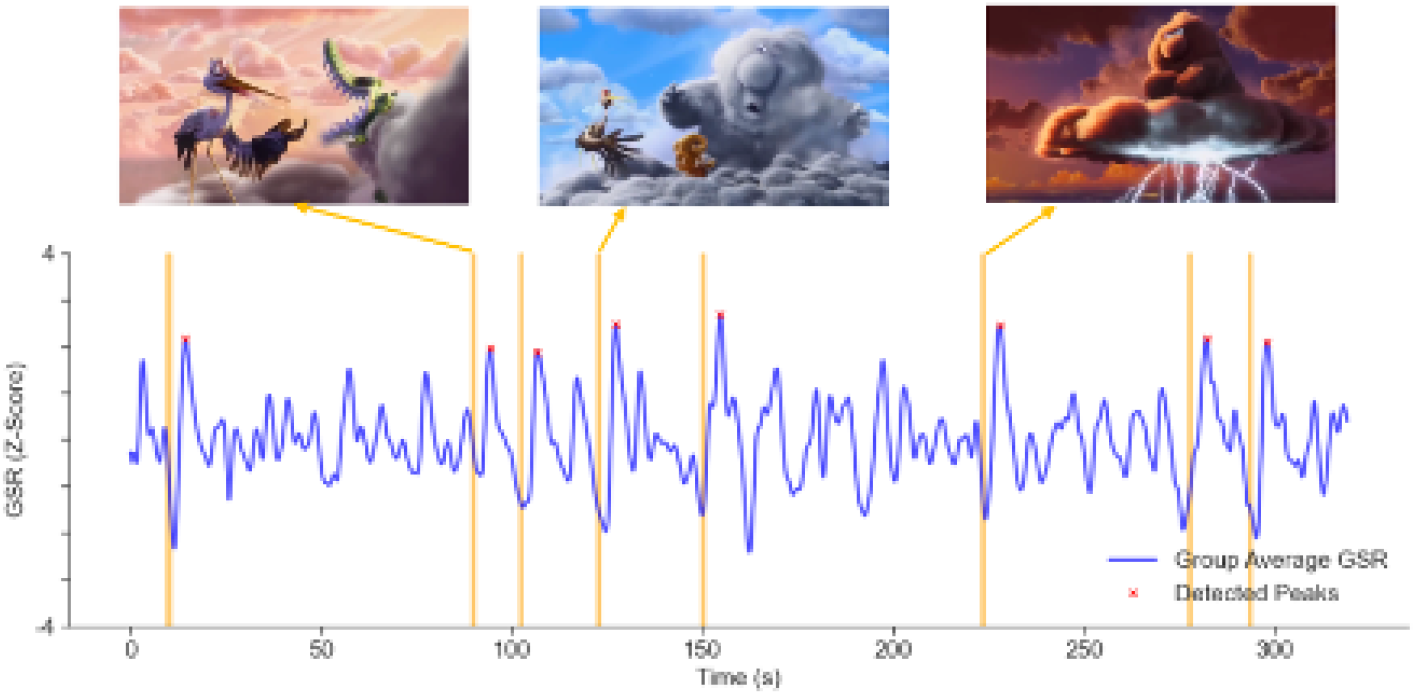
Reverse Correlation Analysis Identifies Highly Salient Movie Scenes. Shown is the group-averaged GSR-trace (main movie) and the top-8 peaks. From these peak times (t(peak)), we went back 4.5 sec to account for the lack between the movie trigger time (t(trigger)) and the GSR response and extracted a 3-second window centered around these trigger times. The resulting eight video snippets were content-analyzed and compared against randomly identified control video snippets. Of note, these peaks also converging audience responses because if people responded idiosyncratically, the responses would average out (see e.g. Figure 2 for split-half consistency).

### S2 Dynamics of Shared Responses

A complementary view of the effect of the branching of storylines is provided in Figure S2, which plots dynamic ISC results, computed separately for each audience via a sliding window analysis (*window length* = 15 sec, *step* = 1). Thus, the blue line represents the moment-to-moment intra-audience correlation for people watching the original movie (main audience), the red line shows the same analysis for people watching the edited version (manipulated audience). As can be seen, these ISC-dynamics unfold almost identically until the branching of the storyline (also see Figure 3 top left panel, which represents an aggregate of these data). This attests also to the reliability of the dynamic ISC analysis itself. After the stories branch, however, the audience alignment within each group continues at a similar level, but the way they evolve varies. This underscores the notion that each movie exerts a “collective grip” (Hasson et al., 2008) on the audience.

**Figure S2.**
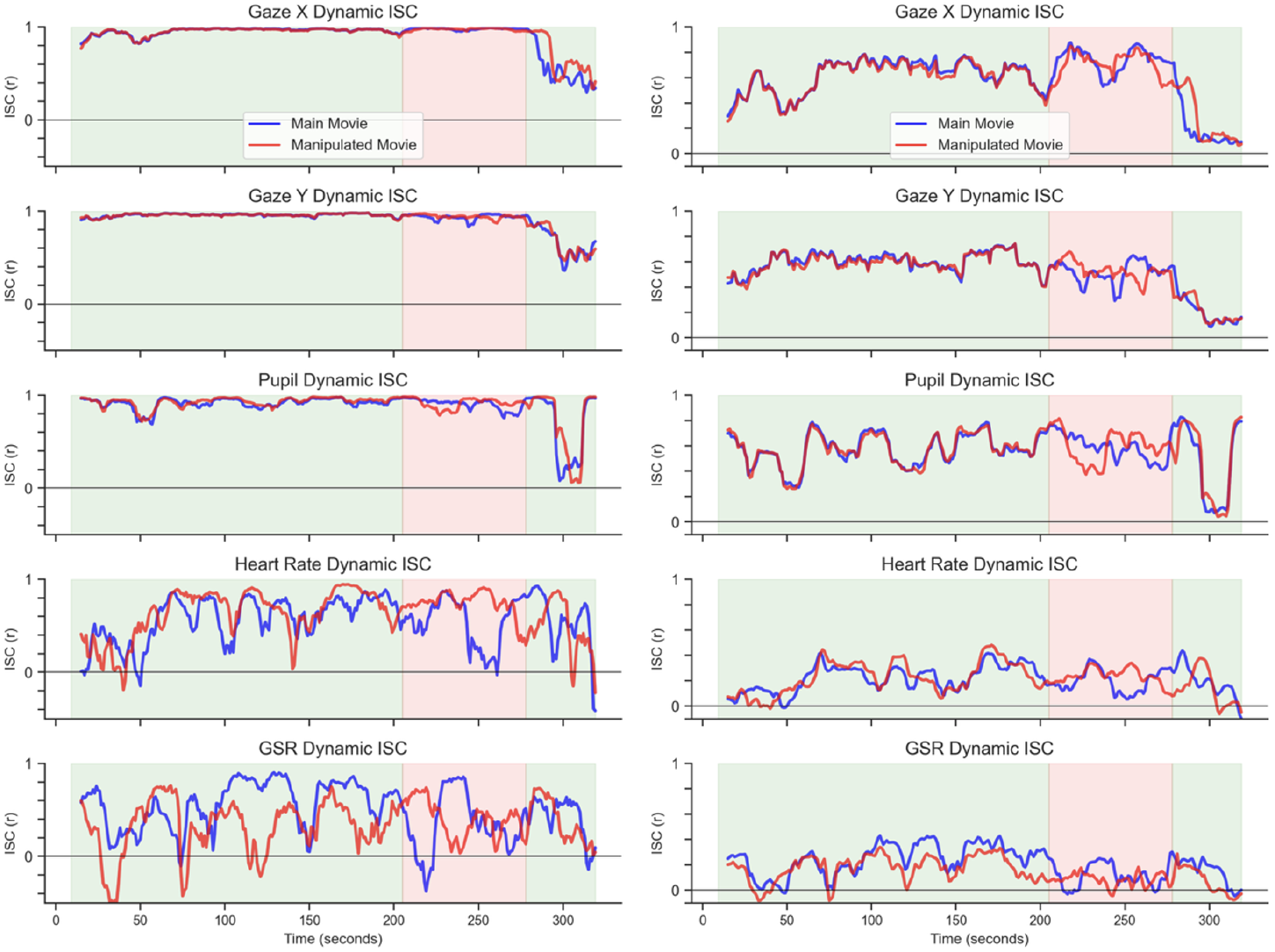
Dynamic ISC (sliding windows) comparing the main and manipulated groups. Results in Figures 2 and 3 are based on analyses of the whole movie or defined segments. This analysis uses a continuous sliding window approach to demonstrate that as long as the input message is the same (until sec 205, when the storyline branches), the collective convergent audience response stays highly similar across all channels, then the collective audience dynamics take a separate trajectory. Left panel: Dynamic ISC for main and manipulated audiences computed using the Split-Half approach. Right panel: Same analysis for the Leave-One-Out approach, showing the same pattern, but at nominally lower level.

### S3 Relationship between Channels: Attentional Alignment Predicts Autonomic Alignment

Another way to look at the hierarchical analysis is to ask whether information acquisition (visual attention) acts as a gateway for downstream autonomic regulation (cf. Kettunen et al., 1998). This issue can be examined by testing whether the fidelity of a participant’s gaze behavior predicts the strength of their physiological coupling with the group. To address this question, we adopted a Leave-One-Out (LOO) Inter-Subject Correlation approach to derive a single “alignment score” per participant for each physiological channel. For a given participant P and physiological signal S (e.g., Gaze or GSR), the alignment score was calculated as the Pearson correlation between a participant’s time-series and the average time-series of all other participants (*N-1*, see Nastase et al., 2019). This yields a vector of *N* scores for each modality, representing how reliably each individual tracked the group response. Statistically, we quantified this relationship by computing the Pearson correlation between the vector of Gaze LOO scores and the vectors of Autonomic LOO scores across participants (*df = N-2*). This approach avoids statistical dependence issues inherent in comparing raw pairwise matrices and tests whether attentional reliability predicts emotional reliability at the subject level.

The matrix reordering analysis revealed a robust link between the sensory intake of the narrative and the resulting autonomic response. As shown in Figure S3 and confirmed by statistical analysis, sorting the sample based on gaze reliability created a progressively weakening gradient.

**Figure S3.**
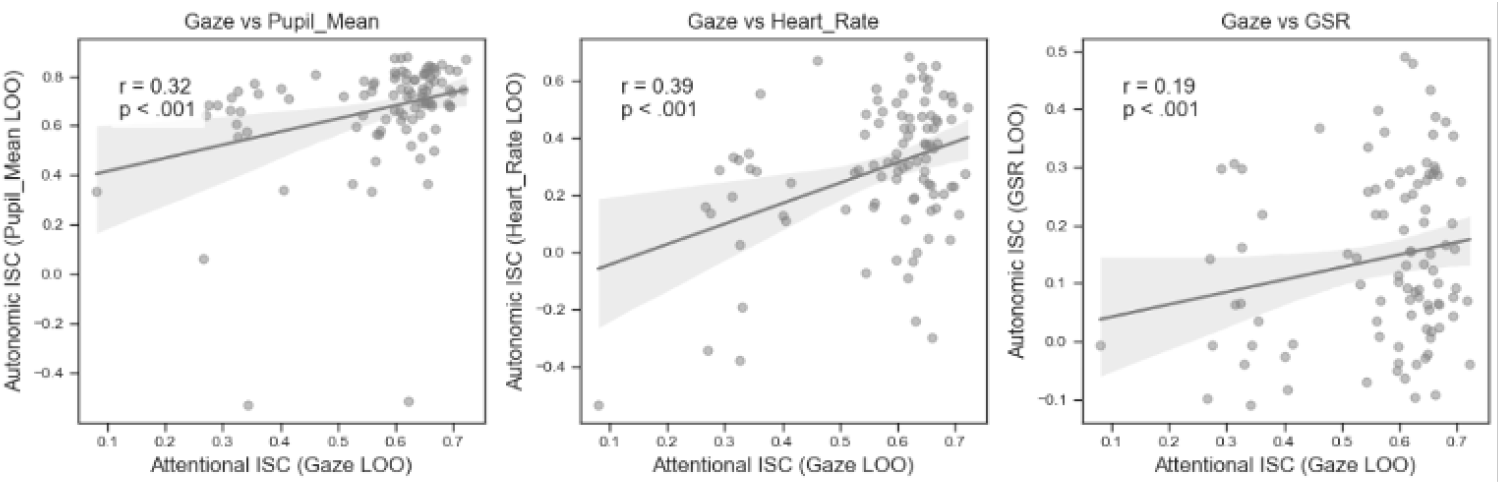
Relationship Between ISC Across Channels. Aggregated LOO-score (by participant) are used to predict Pupil, Heart Rate, and GSR alignment, showing that high Gaze-alignment predicts the other measures.

We found a significant positive correlation between an individual’s Gaze Synchrony and their Heart Rate Synchrony. Thus, people who view the movie in more similar ways tend also to respond more similarly in terms of their heart rate and skin conductance modulations.

### S4 No Effects of Prior Exposure to the Movie

One potential confound for the effects of the movie is prior knowledge. Although we know that even repeated viewing elicits reliable and inter-subjectively correlated responses, it is also conceivable that prior knowledge could affect results since e.g. people would know that there will be a happy ending. This is especially relevant for the manipulated movie, which could additionally trigger expectancy violations and surprise as the story branches. To address this, we asked participants whether they had seen the movie, which 17 percent (17) in the main audience and 20 percent (20) in the manipulated audience answered positively. Thus, we also ran the same analyses excluding these subjects, finding no effects (see Figure S4).

**Figure S4.**
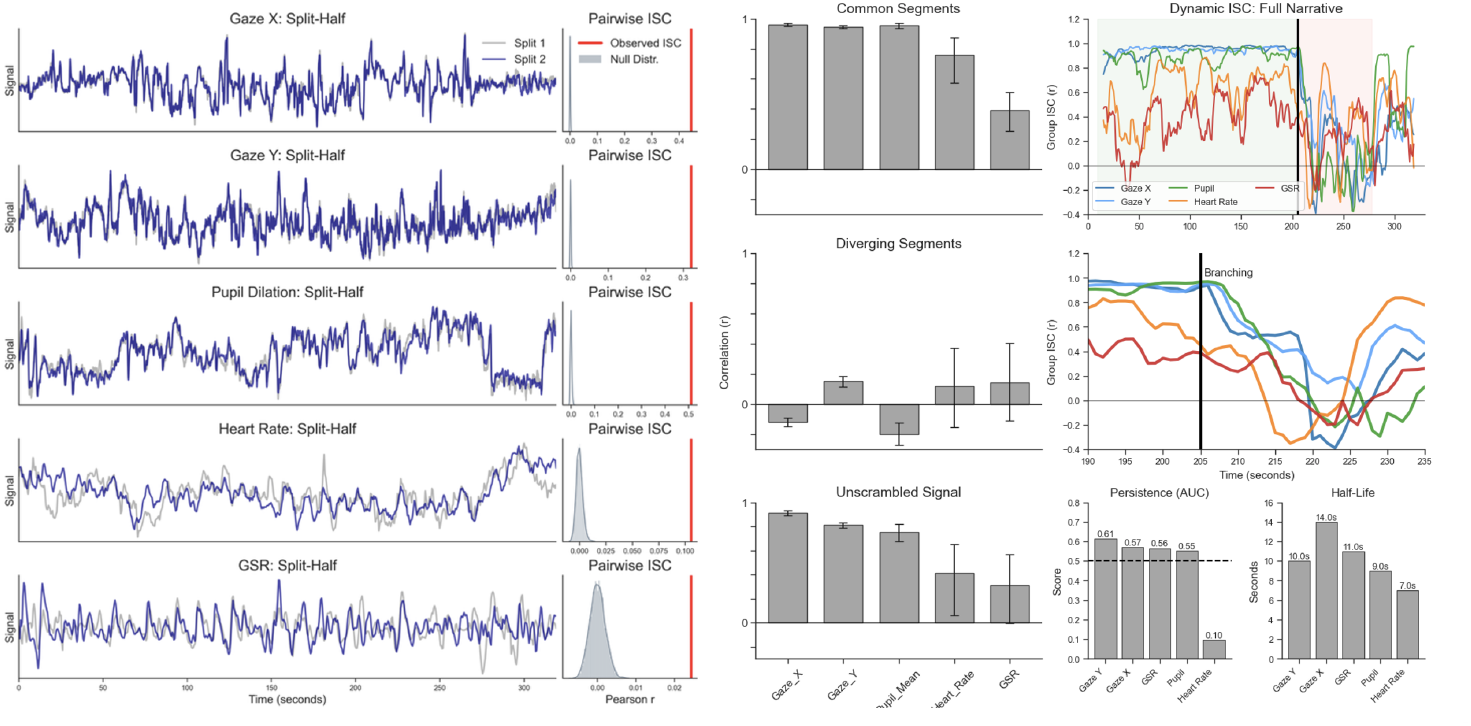
No Prior Exposure Effects. These figures are copies of the plots shown in Figure 2 and 3 in main paper, computed only over data from participants who had not seen the movie before. As can be seen, all hypotheses are fully replicated and no relevant differences emerged.

## Footnotes

2 We caution against conflating alignment with the determinism of “Magic Bullet” or “Hypodermic Needle” models. Identifying the convergent component of reception does not negate individual interpretation; rather, the approach focuses on commonality in responses to the same message, which is necessary for mass communication and central to understanding how it can even occur.

3 Note that this study does not examine brain responses. Previous work has used the same stimulus used here and measured fMRI data (Richardson et al., 2018; Grady et al., 2022), but this article focuses on physiological responses. Including neuroimaging would go beyond the scope of this article and also face some methodological hurdles like varying sample size and presentation conditions, although it is possible and promising as an avenue for future research.

